# GTP Hydrolysis by eIF5B in the Last Step of Translation Initiation Is Activated by a Rotation of the Small Ribosomal Subunit

**DOI:** 10.1101/172825

**Authors:** Israel S. Fernández, V. Ramakrishnan

## Abstract

Placement of an initiator aminoacyl-tRNA [(f)Met-tRNA_i_^(f)Met^] base paired with the AUG initiation codon of a messenger RNA (mRNA) is the first step of translation. The eukaryotic translation factor eIF5B or its bacerial homologue IF2 facilitate the correct positioning of initiator tRNA in the P site of the ribosome. We report the electron cryomicroscopy (cryoEM) structure of a stabilized intermediate state of a yeast 80S/tRNAi^Met^/eIF5B complex at 3.6 Å resolution. The structure shows how a universally conserved tyrosine couples the rotational state of the small ribosomal subunit with GTP hydrolysis.

---

Initiator aminoacyl-tRNA (fMet-tRNA_i_^fMet^ in Bacteria, Met-tRNA_i_^Met^ in Archaea and Eukarya) is the only aminoacyl-tRNA delivered to the peptidyl site (P site) of the ribosome during the initiation stage of translation [1]. Since initiation is not only a rate-limiting step but also of fundamental importance in setting up the correct reading frame, the molecular mechanisms involved in this step are complex and tightly regulated.

The detailed process varies significantly in eukaryotes, where the multisubunit factor eIF4F binds to the 5’ cap of mRNA and recruites the 43S complex of the small subunit (40S) with factors eIF3, eIF2, eIF1, eIF1A, and eIF5 [2, 3].This 48S complex scans along the mRNA until the start codon is reached. This results in conformational changes that lead to GTP hydrolysis by eIF2 and dissociation of the various initiation factors [4]. In the final step, binding of the GTPase eIF5B results in recruitment of the large subunit (60S), followed by GTP hydrolysis and dissociation of eIF5B [1].

Just three of the eukaryotic initiation factors have orthologs in bacteria, and recent work suggests that certain aspects of the mechanism involving these three factors are conserved in both domains of life [5].In particular, the final step involving recruitment of the large subunit by IF2, the bacterial counterpart of eIF5B, is probably the most similar. However, the considerable biochemical and structural data for the bacterial case contrasts with the relatively poor information for the eukaryotic case.

Structures of both archaeal aIF2 [6] and eukaryotic eIF5B [7]have been solved in isolation. Recent studies on yeast eIF5B propose a “domain release” mechanism by which domains III and IV of eIF5B (and the α-helix connecting them, h12) change conformation dramatically with respect to the G domain and domain II upon GTP binding [7]. These conformational changes in domains III and IV are important for Met-tRNA_i_^Met^ binding (domain IV[8]) and ribosome interaction (domain III[9]).

We previously reported a cryoEM structure at 6.6 Å resolution of a yeast initiation complex with Met-tRNA_i_^Met^ and eIF5B, stalled with a non-hydrolysable GTP analog [9]. Comparison with a previous cryoEM structure of the bacterial counterpart [10] confirmed a similar semi-rotated conformation of the 40S as well as a significant repositioning of domains III and IV of eIF5B when compared with the reported crystal structure of the archaeal homolog in isolation [6]. In the ribosome stabilized conformation, eIF5B is anchored to the 60S via its G domain and to the 40S through domains II and III, projecting domain IV deep into the ribosome, in the vicinity of the peptidyl transfer center (PTC).

The repositioning of domain III and IV drags the α-helix connecting them (h12) into the intersubunit space so that the tip of the helix is closely packed against the highly conserved sarcin-ricin loop (SRL) of the 25S rRNA, which is involved in activation of translational GTPases[11]. However, the limited resolution of the structure precluded further structural insights. Here, we report a 3.6 Å resolution structure of eIF5B bound to the ribosome with initiator tRNA that allows us to propose a detailed model for the mechanism for this final step of translational initiation.

Using our previously optimized *in-vitro* initiation reaction set-up with purified components[9] as well as novel computational sorting algorithms implemented in RELION[12], we were able to isolate and refine a homogeneous group of 29,712 cryoEM particles reaching a final resolution of 3.6Å (**Fig. S1**).

A single tRNA is present in our reconstruction in a hybrid configuration, with the anticodon stem-loop placed at the P site of the 40S and the CCA end stabilized by interactions with the L1 stalk, in an intermediate position between the P and E sites of the 60S (**Fig. S2,a**). Following previous convention[13], we refer to this tRNA conformation as p/PE. The Met-tRNA_i_^Met^ initially added to the reaction has suffered a de-acylation event after its delivery to the P site of the ribosome, and is thus trapped in a hybrid p/PE configuration by the L1 stalk, while the use of a non-hydrolizable GTP prevented dissociation of eIF5B from the ribosome, allowing the 40S to back-rotate to a half-rotated state (**Fig. 1 a,b**). Despite the fact that the tRNA is not in the P/I state interacting via its CCA end with eIF5B, as it would be if it were still acylated, the degree of relative rotation of the subunits and the overall conformation of eIF5B, is in agreement with both our previous lower resolution reconstruction[9] as well as with the recent reconstruction of the bacterial counterpart initiation complex with IF2[14]. We therefore conclude that the conformation observed here for eIF5B represents that during initiation just before GTP hydrolysis.

**Figure 1.**
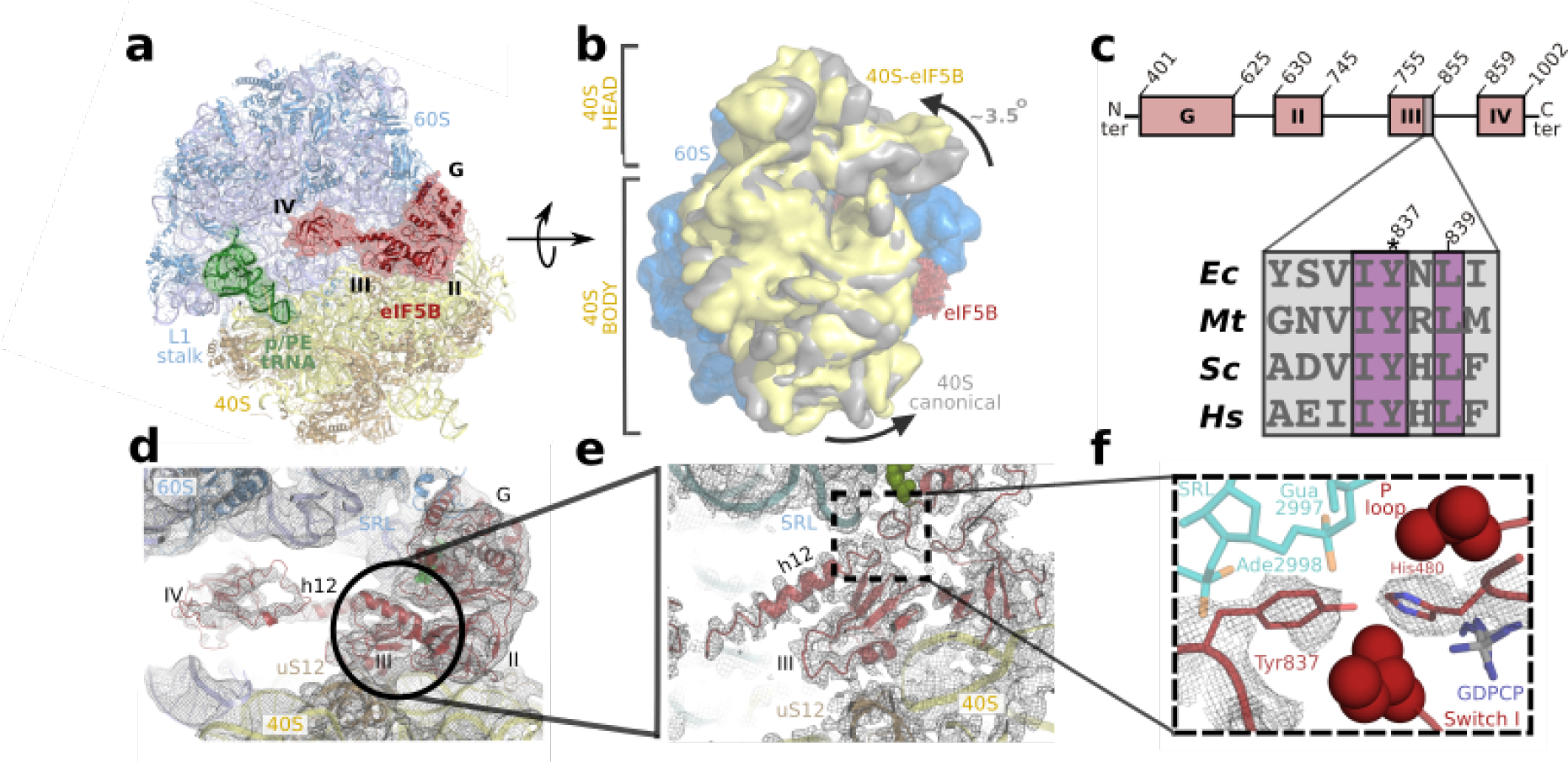
The role of domain III of eIF5B in GTP hydrolysis. (a)eIF5B (red), p/PE tRNA (green) bound to the intersubunit space of the ribosome between the 60S subunit (cyan) and the 40S subunit (yellow) Global conformational changes in the 80S ribosome upon eIF5B binding as viewed from the perspective of the 40S subunit. (**c**) Schematic view of eIF5B domain composition (top), highlighting the position of the conserved segment between domain III and α-helix h12. Bottom, a sequence alignment for this region from Bacteria (*Escherichia coli*, Ec), Archaea (*Methanothermobacter thermautotrophicus*, Mt) and Eukarya (*Saccharomyces cerevisiae*, Sc and *Homo sapiens*, Hs). (**d**) Final unsharpened cryoEM map of ribosome bound eIF5B. Density for the four domains was visible in the final maps. (**e**) Final, post-processed map at 3.6Å focused on domain III of eIF5B (red). Side-chain densities were clearly visible for residues involved in the interaction with the SRL (cyan) as well as for the GTP analog (green Van der Walls sphere representation). (**f**) Detail of the density obtained for the conserved tyrosine residue (Tyr837) in interacting distance with the catalytic histidine residue.

In the current structure, domain III of eIF5B could be observed in the same position as in our lower resolution reconstruction, stabilized between the 40S ribosomal protein uS12 at the bottom and the SRL at the top (**Fig. 1d,e**). Clear densities were seen for the side chains of residues connecting domain III and h12 (**Fig. 1e,f**).

A universally conserved histidine residue in translational GTPases is responsible for the catalytic activation of this family of proteins upon binding to the ribosome[11]. The SRL contributes specific elements that alter the conformation of this key histidine so it is stabilized in a pocket flanked by residues of the switch I and the P loop from the G domain of the translational GTPase. In this activated conformation, the histidine is seen to coordinate a water molecule that is in position for hydrolysis of the γ-phosphate of GTP[11]. This mechanism for GTPase activity seems to be conserved from Bacteria to Eukarya as several high resolution cryoEM structures of ribosome-GTPase complexes from yeast as well as mammals agree well with the initial model derived from X-ray crystallography of a bacterial complex[15, 16]. In the case of eIF5B, the equivalent histidine (His480, *S.cerevisiae* numbering) is stabilized in a similar activated conformation. His480 is further stabilized by interaction with a highly conserved tyrosine residue (Tyr837, **Fig. 1c,f**).

The position and orientation of this conserved tyrosine is determined by the relative rotation of the 40S. A similar configuration has been recently observed in a high resolution cryoEM study for the bacterial complex with fMet-tRNA^fMet^, IF2 and a non-hydrolysable GTP analog[14]. Two mayor populations could be resolved in the bacterial dataset, corresponding to two different rotational sates of the small ribosomal subunit (30S).Thus, in the most rotated structure of the bacterial complex (termed ICI), the bacterial equivalent of Tyr837 is modeled in a position not aligned with the catalytic histidine (**Fig. 2c**, second panel from the left) and the histidine itself is not in the fully activated conformation seen here. The second structure present in the bacterial complex (termed ICII), exhibits an intermediate degree of small subunit rotation similar to the one described here, but unfortunately, the limited local resolution of this complex in that area meant that there was no description of the particular tyrosine[14]. Given the conservation of the factor and the tyrosine in particular(**Fig. 1,c**), we suggest that the structure of the activated state in bacteria is probably very similar to that seen here.

The present structure suggests how the activation of translational GTPases is used to regulate a key step of initiation (**Fig. 2**). Initially, the exchange of GDP to GTP induces a conformational change of switch I and P-loop around the γ-phosphate which triggers the reorganizations of domains III and IV (**Fig. 2a**, left and **b**, “domain release”[7]). The new conformation of domains III and IV facilitates binding of the initiator aminoacyl-tRNA in the context of the small ribosomal subunit (**Fig. 2a**, second panel from the left and **c**, left panel). The subsequent recruitment of the 60S is facilitated by a fully rotated state of the 40S. In this conformation, the catalytic histidine is in a partially activated conformation, due to the conserved tyrosine is not in a position to interact with and stabilize the catalytic histidine in its fully inserted position (**Fig. 2a** central panel, **c**, second panel from the left). The half back-rotation movement of the 40S is transmitted through the ribosomal protein uS12 to the bottom of domain III, which eventually rotates to align the conserved tyrosine with the catalytic histidine, thus stabilizing it in its final activated conformation (**Fig. 2,a** second panel from the right, **c**, second panel from the right). GTP hydrolysis would result in a disordering of switch I, leading eventually to conformational changes on eIF5B and dissociation of the factor from the ribosome. Finally, once the factor has left the ribosome, the 40S can relax to its non-rotated configuration, allowing the initiator aminoacyl-tRNA to occupy a canonical conformation in the P site, with a vacant A site ready for the first round of elongation (**Fig. 2a**, right panel, **c**, right panel).

**Figure 2.**
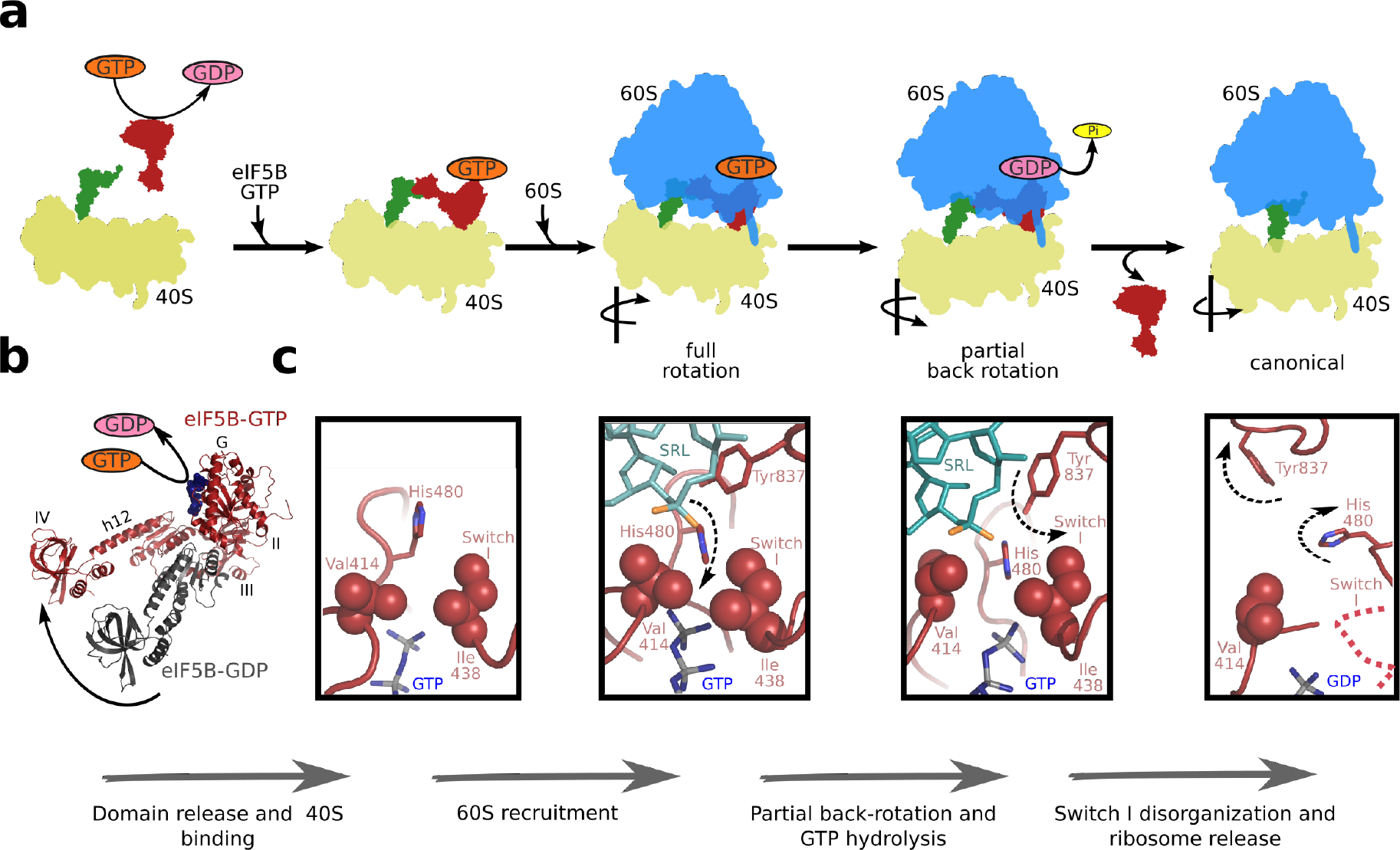
A comprehensive model for large subunit recruitment and eIF5B function. Cartoon representation of the main molecular events proposed for eIF5B mode of action. Starting from the left: interchange of GDP (pink) to GTP (orange) allows for the recognition of the initiator aminoacyl-tRNA (green) by eIF5B (red) in the context of the smal ribosomal subunit (40S, yellow). Recruitment of the large ribosomal subunit (60S, blue) is facilitated by eIF5B-GTP in the fully rotated state of the 40S[14]. Partial back-rotation of the 40S results in interaction of a universally conserved tyrosine and the active site histidine, which results in a catalytically active form for GTP hydrolysis. After GTP hydrolysis and phosphate release, conformational changes in eIF5B-GDP result in its dissociation from the ribosome and a full back-rotation of the 40S, to a non-rotated configuration of the ribosome with initiator aminoacyl-tRNA in a canonical P/P state. (**b**) Superposition of the eIF5B-GDP (PDB-ID 4N3N[7]) and eIF5B-GTP (from this study), showing the re-orientation of domains III and IV and the linker α-helix h12. (**c**) Diagram of sequence of events described in (**a**) showing the region around the γ-phosphate of GTP (blue). Adapted from PDB-IDs 4NCN, 4NCF[7] and 3JCJ, 3JCN [14]. This sequence of events is suggested by a common activation mechanism for translational GTPases implied by structural data.

The structure shows how the dynamical properties of both the ribosome and eIF5B are used to regulate the final step of initiation. In this model, GTP hydrolysis by eIF5B is possible only with the correct conformation of eIF5B and the ribosome, because it is only in that conformation that the universally conserved tyrosine and histidine can interact to stabilize the latter in a conformation that seems to be required for the activation of translational GTPases. Thus, GTP hydrolysis by eIF5B serves as a check point for entering the elongation cycle, that ensures appropriate binding of initiator tRNA and the large ribosomal subunit.

## METHODS

### Preparation of ribosomes and initiation factors

Ribosomal subunits were purified from *Kluiveromyces lactis* (strain GG799) and used in an *in-vitro* initiation reaction as previously described[9], except that eIF5B was from *K.lactis* (see below) and the final buffer was 10 mM 2-(N-morpholino)ethanesulfonic acid (MES) adjusted to a pH of 6.5 with potassium hydroxide.

### *K.lactis* eIF5B production

An N-terminally truncated eIF5B version (residues 396-1002, with a molecular weight of 68 kDa) was cloned from *K.lactis* genomic DNA in a T7 based expression vector that added a 6xHis-tag followed by a TEV protease cleavage site at the N-terminal. Over-expression in *E.coli* strain BL21-AI (Invitrogen) was followed by chromatography using HisTrap, Q-HP and Sephacryl S-200 (GE Healthcare). The purified protein was concentrated to 100 *µ*M and snap frozen in liquid nitrogen.

### tRNA and mRNA production

Yeast Met-tRNA_i_^Met^ was purified as previously described[9].The mRNA with sequence 5’GGAA[UC]_4_UAUG[CU]_4_C_3_ was chemically synthesized by IDT.

### Electron microscopy

Aliquots of 3*µ*l of the 80S initiation complex at a concentration of 80 nM were incubated for 30s on glow-discharged holey carbon grids (Quantifoil R2/2), on which a home-made continuous thin carbon film (estimated to be 30Å thick) had previously been deposited. Grids were blotted for 2.5s and flash cooled in liquid ethane using an FEI Vitrobot. Grids were transferred to an FEI Polara G2 microscope that was operated at 300 kV. Defocus values in the final data set ranged from 1.6-3.6 *µ*m. Images were recorded using the FEI software EPU on a back-thinned FEI Falcon III detector at a pixel size of 1.07Å. The individual frames from the detector (36 frames for each 1s exposure) were captured and stored on disk. All electron micrographs were evaluated for astigmatism and drift.

### Image processing

CTF estimation was performed with GCTF[17] and automatic particle picking was done with Gautomatch without a specific template using a diameter of 280 pixels. All 2D and 3D refinements were performed using RELION [18]. We used reference-free 2D class averaging to discard 60S subunits and defective particles, resulting in 64,815 particles of the final data set for subsequent 3D refinement and classification. Refinement of all particles against a single model (a 60Å low-pass filtered version of EMDB-2275) yielded a preliminary, consensus reconstruction with local fuzzy density for the 40S subunit, the L1 stalk and eIF5B. Subsequently, we employed local classification with a masks covering the intersubunit space and signal subtraction[12] to identify a class of 29,712 particles, for which all four domains of eIF5B showed clear density. Reported resolutions are based on the gold-standard FSC=0.143 criterion[19]. Prior to visualization, all density maps were corrected for the modulation transfer function (MTF) of the detector, and then sharpened by applying a negative B factor that was estimated using automated procedures[20]. Manual rebuilding was done using COOT[21] and the final model was refined with REFMAC[22] following established protocols.

**Figure S1.**
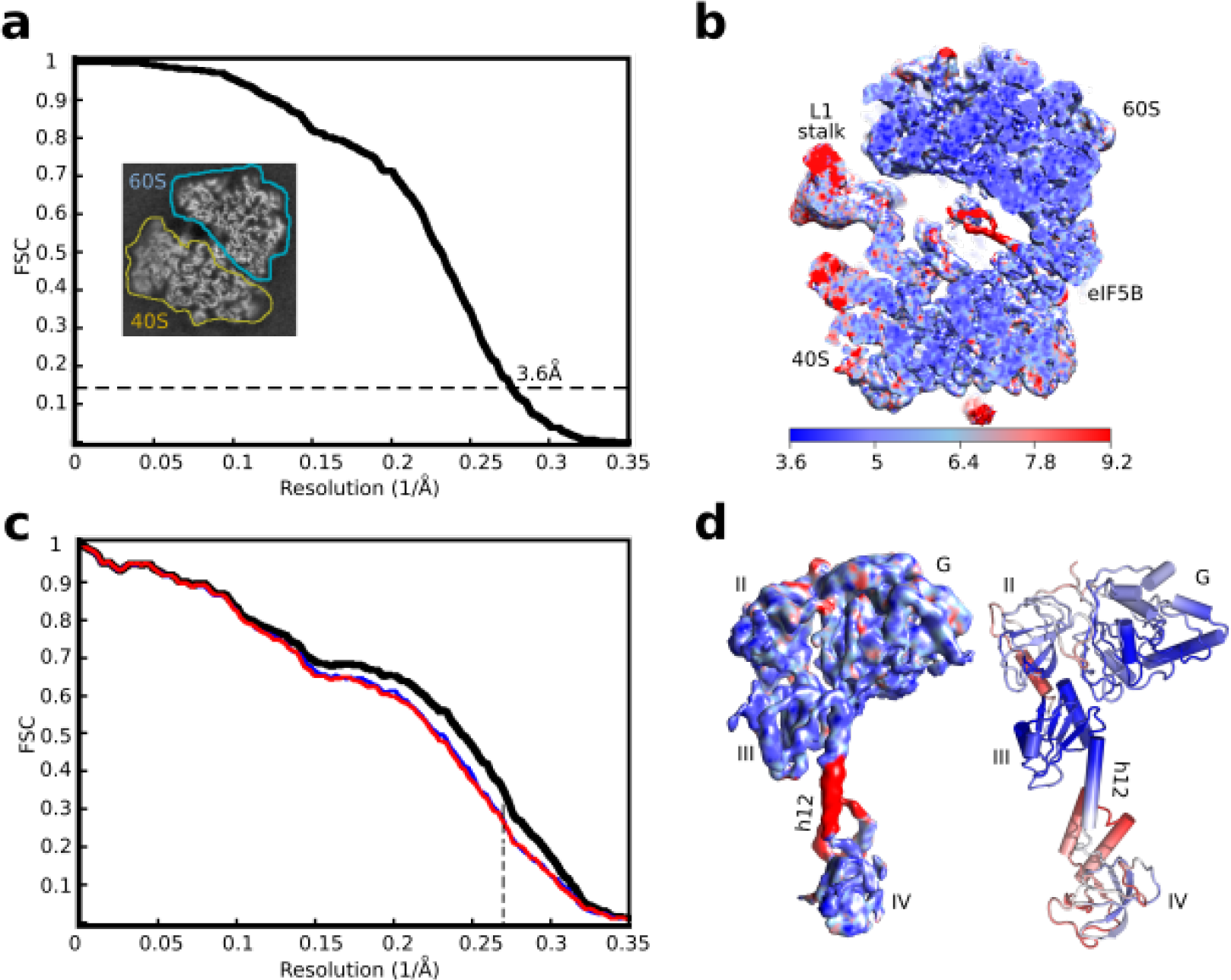
Fourier Shell Correlation curves and local resolution. (**a**) Fourier Shell Correlation (FSC) computed for the two half maps of the final subset of particles after classification. The resolution is estimated to be 3.6Å using the 0.143 criterion[19]. Inset, slice through the final map showing the high local resolution for both ribosomal subunits except for the tip of the head of the 40S, which appears to be more mobile. (**b**) Slice through the final, unsharpened map colored according to the local resolution as reported by RESMAP[23]. Resolution beyond 4Å can be observed for the 60S and 40S as well as for domains G,II and III of eIF5B. More flexible areas like the L1-stalk, the tip of the head of the 40S and domain IV of eIF5B are observed at lower resolution. (**c**) Map-versus-model cross validation FSC. The final model was validated using standard procedures. FSC of the refined model against half map 1 (blue) overlaps with the FSC against half map 2 (red, not included in the refinement). The black curve corresponds to the FSC of the final model against the final map. (**d**) Left, unsharpened density for eIF5B colored according to local resolution using the same scale as in **b**. Right, final model for eIF5B colored according to the refined atomic B-factors computed by REFMAC[22] (blue low, red high).

**Figure S2.**
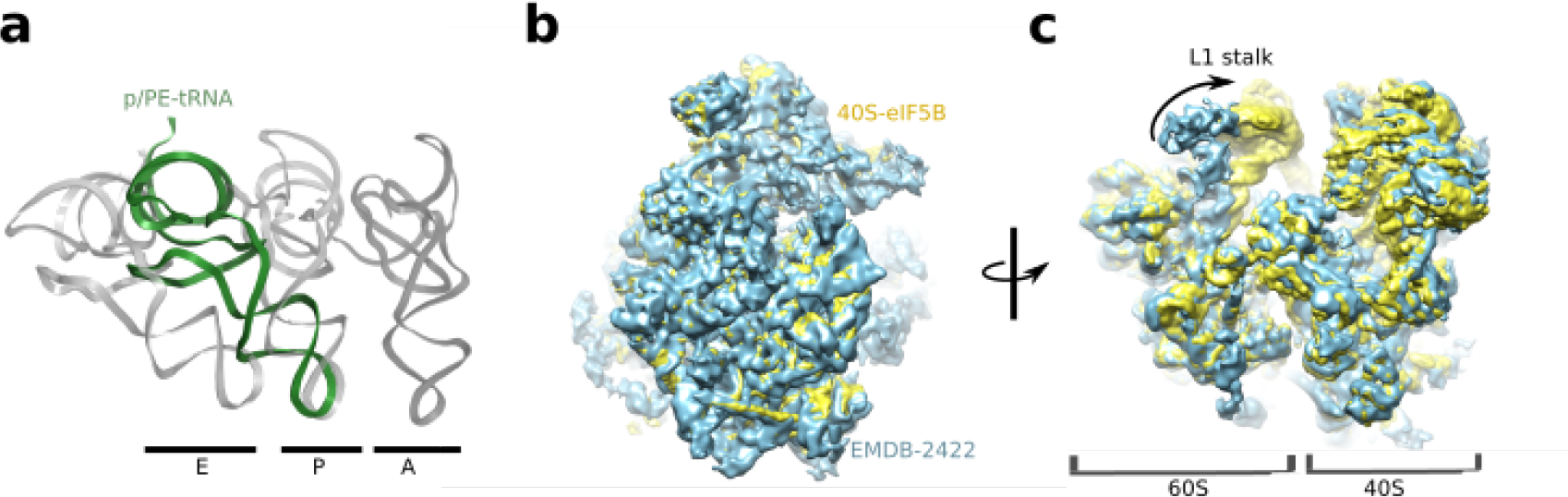
tRNA conformation and superposition with low resolution map. (**a**) Final refined model for the tRNA in the present reconstruction (green) superposed with canonical tRNAs in grey (PDB-ID 4V5C[24]). The anticodon stem-loop is located in the P site of the 40S whereas the CCA end is placed in between the P and E sites of the 60S (p/PE configuration). (**b**) Superposition of our previous lower-resolution reconstruction (cyan, EMDB-2422[9]) with the present one (yellow) showing the similar degree of 40S rotation.(**c**) A 90 degree rotated view from the position showed in (**b**). The arrow highlights an inward displacement of the L1-stalk in the current structure that stabilize the tRNA.

## Acknowledgements

We are grateful to Christos Savva for technical support with cryo-EM, Greg McMullan for help in movie data acquisition, Toby Darling and Jake Grimmett for help with computing and Sjors Scheres for advice in image processing. VR is funded by grants from the UK Medical Research Council (MC U105184332), the Wellcome Trust (WT096570), the Agouron Institute and the Louis-Jeantet Foundation. Electron microscopy maps and coordinate files have been deposited in the EMDB with accession numbers XXXX and YYYY.

